# Scattering-driven PPG signal model

**DOI:** 10.1101/2021.06.22.449402

**Authors:** I. Fine, A. Kaminsky

## Abstract

This article discusses the origin of photoplethysmographic (PPG) signals. Two plausible hypotheses that could explain the phenomenon of pulsatile optical signals are analyzed: the volumetric hypothesis in which changes in the arterial blood volume are responsible for the observed signal pulsation, and a model wherein changes in the measured signal are driven by the size fluctuations of red blood cell (RBC) aggregates. The theoretical approximation where the size of scattering particles representing RBC aggregates varies as a function of pulsatile changes in blood flow is elaborated. Within this model’s framework, the gamma coefficient used in pulse oximetry was calculated for the volumetric-related and aggregation-related models. Two pairs of wavelengths, (670 nm, 940 nm) and (590 nm, 940 nm), were selected to determine gamma. As a function of aggregate size, the gamma behavior was simulated for these pairs and the two hypotheses. To verify the model predictions experimentally, the PPG signals at the fingertip were measured using reflection geometry. Two combinations of light-emitting diodes with two pairs of wavelengths were utilized as light sources. To manipulate the length of aggregates in the blood, external pressure was applied to the fingertip, presumably reducing the blood flow velocity. The gamma values were determined. The derived results fully agree with the theoretical predictions of the aggregation-driven PPG signal model. In addition, using a pressure sensor, the oscillometric signal amplitude in the fingertip and the PPG signal amplitude were simultaneously measured. The comparison results of oscillometric and optical signals at elevated external pressure values are not consistent with the volumetric hypothesis. All of the foregoing experimental results strongly support the argument favoring the incorporation of the proposed aggregation mechanism into the generic PPG signal model.

## Introduction

The pulse oximeter is one of the most widely used medical devices. The operating principle of pulse-oximetry is based on the fact that the light intensity transmitted or reflected from a finger or an earlobe varies with the pulse wave [1]. This type of measurement is called photoplethysmography (PPG). The widely accepted assumption describes the PPG signal in terms of pulsatile changes in the vascular blood volume [2]. The fact that pulsation changes in the blood volume are measurable by various methods cannot be disputed; however, this does not mean that only volumetric changes are responsible for the pulsatile appearance of optical signals. In 1967, the question pertaining to the origin of PPG signals was raised for the first time when it was experimentally shown [3] that optical pulsatile components appeared in the absence of measurable volumetric changes. In 2013, a published article reported that a considerable quantity of pulsating optical signals was detected in various human bones although the number of red blood cells during the cardiac cycles remained constant [4]; nevertheless, these findings were barely discussed in the scientific literature [6]. In 2000, to explain the optical signal behavior *in vivo*, accounting for the RBC aggregation process or rouleau formation was proposed [5]. The same study considered the process as responsible for the increase in light transmission during aggregation and blood flow cessation *in vivo*. Note that the relationship between the RBC aggregation and optical signal has been well known and studied for many years *in vitro* [7, 8, 9,10]. Thus, the study in [5] assumed that the manifestation of the dependence of reflected or transmitted light on the state of aggregation of RBCs is not limited under *in vitro* conditions but also *in vivo* [11]. Under *in vitro* and *in vivo* conditions, the presence and magnitude of shear forces in blood flow must be attributed to the key elements regulating the rouleau size. In subsequent years, a specific mechanism explicitly pertaining to the effect of aggregation on the optical signal was quantitatively elaborated [12]. The qualitative understanding of the proposed model can be summarized as follows. The dynamic balance between the destruction and formation of RBC aggregates is determined by the blood velocity. The shear rate is defined as the slope of the velocity profile. This rate is high when the flow velocity is high, and the vessel diameter is small; conversely, the shear rate is low when the flow velocity is low, and the vessel diameter is large. The size of RBC aggregates is inversely proportional to the magnitude of the shear rate [29]. The increase in shear rate results in the breakup of large aggregates into smaller ones or into single RBCs. For shear rates exceeding 50 s−1, all RBC aggregates are separated into single erythrocytes [31]. Under normal conditions, a periodic pressure wave causes oscillations in velocity and shear forces, modulating the aggregate size. Hence, the fluctuation in the shear forces of blood flow is the main rheological factor leading to observed variations in the measured optical signals. The influence of shear forces on the size of aggregates *in vitro* and the semi-empirical description of this process are published elsewhere [13, 14].

To describe the effect of erythrocyte aggregation on the transmission of light, two levels of modeling have to be considered. First, the change in light scattering should be expressed in terms of aggregate size. In [15], the Wentzel–Kramers–Brillouin (WKB) approximation was proposed to describe the light scattering coefficients for long spheroid particles. This method is the approximation of anomalous diffraction and applies to the scatterer of any arbitrary shape and size. The WKB-based solution allows the derivation of a microscopic scattering coefficient for substantially large aggregates. Following this approximation, radiation propagates inside the particle in the same direction as the propagation of incident radiation (low-refractive particles). The radiation wavenumber inside the particle is equal to the radiation wavenumber in the particle material. In this case, the effect of scattering is related to the radiation phase change when radiation propagates inside the particle. Based on this model, the erythrocyte aggregate can be approximated as a growing spheroid. This model is suitable for describing a long aggregate that grows due to the blood stasis process. However, in the case of blood flowing through arterioles, the assumption that the average rouleau size can exceed several erythrocytes is difficult to realize; consequently, the elongated spheroid approximation is not that applicable.

For the scattering of light by particles whose sizes are comparable to the wavelength, the Mie model affords an exact analytical solution in the case of spherical particles. Although a single erythrocyte is not spherical, it has become apparent that the Mie solution is well-suited for RBCs [16]. With the transition of one erythrocyte to an aggregate of several erythrocytes, the combined particle resembles a sphere. Therefore, the use of the Mie model in this case is even more justified than in the case of a single RBC. For this reason, approximating the growing small rouleau with an equivalent inflating sphere was proposed [12]. In this way, the change in the scattering cross-section during aggregation can be described by corresponding Mie functions. An additional level of modeling refers to the propagation of light in tissues and blood [17]. A wide variety of models ranging from the transport equations of photon flux in blood to the Monte Carlo models of light propagation in tissue and blood exists [19, 20, 22]. The selection of a suitable model depends on the practical purpose of the model. In this case, the objective is to describe the variable pulsating component of the optical signal, which is completely predetermined by changes in the optical properties of RBCs. The tissue surrounding the blood vessels is responsible for causing the multiple scattering of light; accordingly, this justifies the adoption of diffusion approximation. Because the goal is to describe the variable in this study, experiments are implemented to test the plausibility of the two concepts for the PPG signal: the volumetric model and aggregation model (also named scattering-driven model (SDM)) [12]. The behavior of “gamma” values as a function of the aggregate size of two pairs of wavelengths (for example (670 nm, 940 nm) and (590 nm, 940 nm)) was simulated for both models. Experimentally, the change in the aggregate size was achieved by reducing the blood flow at the measurement site where external pressure was applied. Moreover, volumetric pulsations were recorded by measuring the pressure in the air cushion wrapped around the fingertip (oscillometric signal). These measurements were simultaneously performed with the detection of PPG signals. In a series of experiments, the amplitude and phase shift between the optical and oscillometric signals were investigated under different pressure conditions. The derived experimental data were compared with the model predictions for the two discussed cases.

## Theoretical consideration

### RBC aggregation and shear rate

An RBC can be approximated as an oblate spheroid with an axial ratio of approximately 0.3 and a maximum radius of 4.1 microns. The RBCs in the plasma tend to form aggregates that appear similar to a stack of coins [28]. The so-called rouleau formation is caused by plasma macromolecules and is supposed to be a reversible aggregation process. In flowing blood, an inverse relationship exists between the size of red blood cell aggregates and the shear rate of blood *in vivo* [30].

In the simplified case, when blood flow is laminar and the vessel is cylindrical, the velocity profile, *v*(*r*), can be approximated [31] as

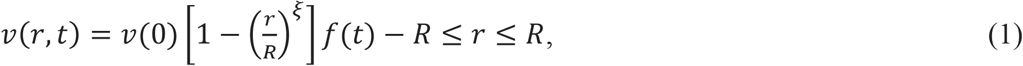

where *v*(0) is the maximum velocity at the center position (r=0); R is the radius of the artery; *f(t)* is a periodic function, which is driven by the difference between systolic and diastolic pressure waves; and ξ, which ranges from 2 to 4 at normal flow rates, represents the degree of blunting (the bluntness factor controls the shape of the velocity distribution, e.g., *ξ* = 2 results in a parabolic velocity profile). It is given by the following:

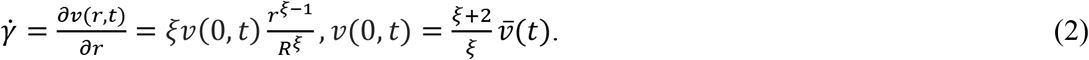

The average value of shear forces within the volume bounded by the radius, R_i_, can be estimated as follows:

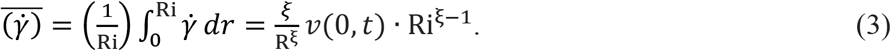

For small arterials [25], the variation in velocity from systolic to diastolic phases is 1.5–2.5 mm/s. The number of red blood cells in the rouleau depends on the shear forces at any given instance. Assume that the rate of aggregate destruction in the systolic part is sufficiently fast for the disaggregation process not to lag behind the pulse velocity wave. For RBCs aggregating in the lateral direction of vessels, the average rouleau size (An) is given by [24]

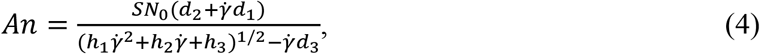

where S=0.6/ϔ, *N*_0_ = 4.96 × 10^−3^ *μ*^−3^, d_1_=3×10^3^ *μ*^3^, d_2_=1.8×10^3^ *μ*^3^, d_3_=1, h_1_=d_3_^2^+2d_1_ d_3_SN_0_−d_1_^2^S^2^N^2^, h_2_=2d_2_d_3_SN_0_−2d_1_d_2_S^2^N_0_^2^, and h_3_=−d_2_^2^S^2^N_0_^2^.

Figure 1(a) shows a simulation of a pulse wave whose average velocity varies in the range 1.5–2.5 mm/s. Based on Equations (1)–(3) and considering a 200-μm thick arterial vessel as an example, the average change in the shear rate caused by velocity fluctuations when Ri =100 μm is 4–5 s^−1^; when Ri=50 μm, the shear rate drops to the extremely low values between 0.5 and 1 s−1 (Figure 1(b)). Using Equation (4), Figure 1(c) demonstrates the variation in length as a function of shear rate. The figure indicates that for small values of shear forces, the extremely small variations in the shear rate considerably affect the aggregate size.

**Figure 1:**
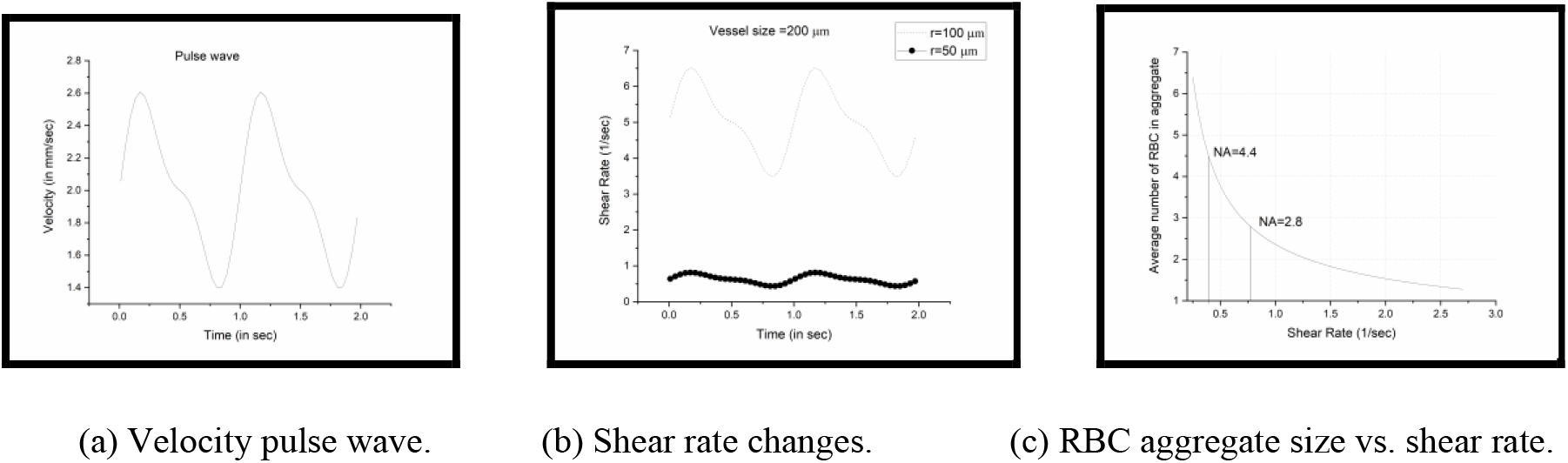

The final signal is virtually formed by averaging the signals over an ensemble of different vessels, where the shear rate distribution is defined by the vessel diameter and distribution of pulse wave velocities. In addition, note that for small arterioles, the degree of blunting approaches 4; this further increases the dependence of aggregate length on the shear forces in the central region of the vessel.

### Gamma determination

The PPG signal is characterized by light intensity changes after passing through a perfused tissue. The associated pressure waves lead to periodic changes in the transmitted and reflected light. The focus of this work is on the specific mechanism that causes changes in the intensity of the measured signal. In pulse oximetry, the signal variations are converted into the “gamma” value, commonly called the “ratio of ratios.” This value is calculated from the pulsatile (alternating current (AC)) and non-pulsatile (direct current (DC)) components of the measured signal for two wavelengths (i.e., λ_1_ and λ_*ref*_) [1]:

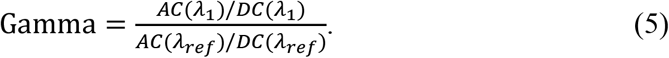

In pulse oximetry, the “gamma” value uniquely determines the percentage of oxyhemoglobin in arterial blood (SPO2). Another parameter, called parametric slope (PS), can be regarded as virtually equivalent to “gamma” [5] and was found to be a more useful form to express “gamma” for model analysis.

Let the intensity of the measured optical signal be denoted by I(λ, t), which is a function of time and wavelength. At any given time, the derivatives of the signal can be calculated. As a rule, the ratio of signal derivatives measured at two wavelengths does not considerably fluctuate at different pulse wave intervals. The average PS value over the full pulse wave interval corresponds to the standard “gamma” (the “ratio of ratios”):

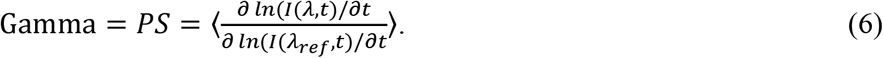

In practice, LEDs are used as light sources in pulse oximetry. In this case, the spectral intensity distributions, W_1_(λ) and W_2_(λ), for each LED must be considered, and Equation (6) must be modified, as follows:

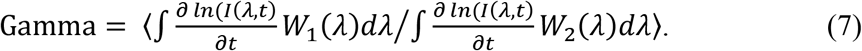

The signal intensity, I(λ, *t*), passing through or reflecting from the tissue depends on the optical properties of the tissue; however, temporal pulsation dependence is determined only by the blood component. Accordingly, interest is focused on the time-dependent function of the scattering and absorption coefficients of blood. These coefficients can be expressed in terms of the scattering and absorption cross-sections [15]:

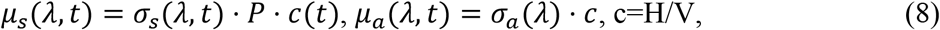

where c is the concentration of scatterers; H is the hematocrit value defined by the relative volume occupied by RBCs; σ_*a*_(λ) is the absorption cross-section of RBCs; σ_*s*_(λ, *t*) is the total scattering cross-section of the scatterer; V(t) is the mean volume of scatterer; and *P* is the packing factor (initially introduced by Twersky [18] and then experimentally adjusted to blood). For the suspension of single RBCs, this factor is commonly taken as *P* = *H*(1.4 − *H*). The concentration of scatterers, c(t), each consisting of N_R_ erythrocytes, is given by c(t)= H/(*V*_0_ ∗ *N*_*R*_).

The absorption cross-section of blood depends on the hemoglobin saturation given by

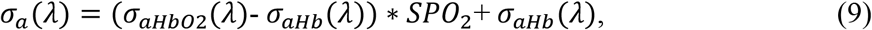

where σ_*aHbO*_2__ and σ_*aHb*_ are the absorption cross-sections for oxyhemoglobin and deoxyhemoglobin, respectively; SPO_2_=HbO_2_/([HbO_2_]+[Hb]) (here, HbO_2_ and Hb are the concentrations of oxyhemoglobin and deoxyhemoglobin, respectively).

The light scattering by a single erythrocyte is well described by the scattering function of Mie [16], which provides a complete solution for determining the scattering cross-section value and phase-scattering function (g). To adjust the Mie theory for aggregates, the following simplified model is used. When RBCs stick together in a chain, the new scattering particle can be approximated as a sphere (Figure 2). As the number of erythrocytes participating in aggregation approaches 4μ, the shape of the resulting particle becomes more symmetric and more closely corresponds to the approximation of a sphere.

**Figure 2:**
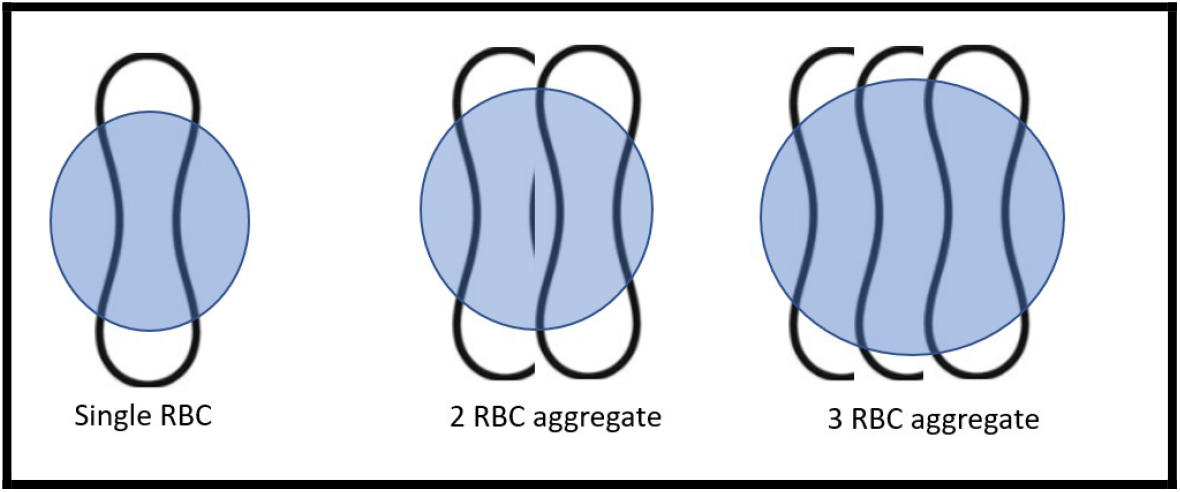
Rouleaux approximated by equivalent sphere.

The particle is considered composed of NR erythrocytes (each with volume *V*_0_). For a given size of small aggregates, an equivalent sphere with radius reff generating the same volume as the particle composed of NR erythrocytes is produced. The effective radius, reff, of each aggregate is given by

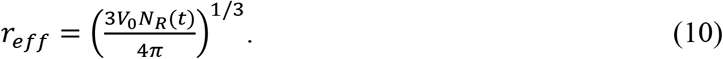

A diffusion model [20] is typically employed to describe the transmission of light through tissue. The “inverse diffusion length” or diffusion coefficient is convenient to use as a basic parameter. This coefficient [26] is defined by

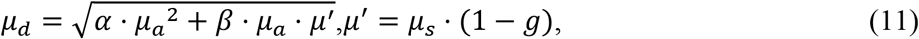

where g is the scattering anisotropy factor, which is dependent on the size of the scattering particle and wavelength; *α* and *β* are adjustable parameters (taken as 1 based on further analysis).

According to the Mie model, the scattering cross-section and g are defined by the radius of the sphere, r_eff_, and relative refractive index. It was shown [27] that an ellipsoid with a revolution of a given size, axial ratio, and orientation could be approximated by a suitable equivalent sphere. The effects of the difference between the ellipsoid and equivalent sphere can be accounted for using a corrected refractive index, m_c_, for the latter [27]:

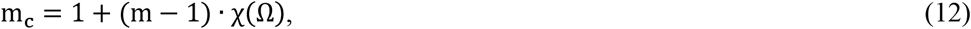

where m is the refractive index of the particle relative to the surroundings; χ(Ω) can be expressed in terms of the axial ratio parameters of the ellipsoid and angle (Ω) of incidence of light relative to the main axis of the ellipsoid. In the rouleau analysis, the foregoing factor was considered in calculating the scattering cross-section and g.

Considering the second level of modeling, a diffusion equation suitable for the objectives had to be selected. To assess the dependence of the results of the calculated gamma on the type of diffusion model, two approaches focusing on the diffusion of light in a highly scattering medium were selected. Because the measuring system is designed for the reflective signal, a model in which the intensity of the measured reflected signal is expressed in terms of the absorption scattering and diffusion coefficient is selected [21]:

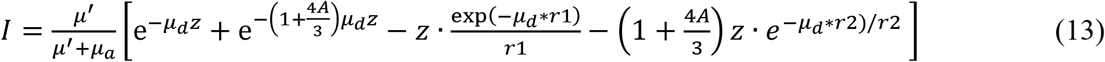

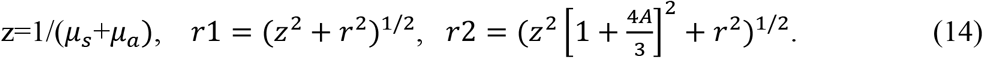

Parameter A depends on the refractive index of the medium (A =1 for n =1, and A > 1 for n > 1); r is the detector radius (r= 3 mm in this work).

Within the framework of this model, any changes in the properties of the medium can be expressed through the scattering and absorption coefficients. However, this model is not well-suited for simulating volumetric changes. Accordingly, as a second choice, a simple approximation suitable for both transmission and reflection is applied. Note that the scattering of light in tissue is considerably more isotropic than scattering using erythrocytes; consequently, the photon flux becomes virtually isotropic from the beginning. Thus, according to the simplified diffusion model, the spatial distribution of the diffusing photons is dictated by the properties of the tissue, whereas the hemoglobin in blood particles absorbs the photons. Both transmission and reflection modes can be considered as transillumination governed by diffusion, where the direction of the flux of photons does not play a significant role. The evaluation of signal change due to fluctuations in absorption and scattering caused by the blood component only is of interest. The simple exponential dependence of the diffused signal on the properties of blood is the experimental fact on which pulse oximetry is based. This is the reason for choosing a simple exponential expression of photon diffusion. Only the time-dependent behavior of the signal is relevant. Thus, the following expression governs the changes in the light intensity:

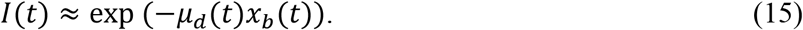

Equation (15) ascribes the diffusion coefficient and volumetric changes to the time-dependent intensity modulation. Let time-dependency be assigned only to x_b_(t); this specific case is called the volumetric model. According to this model, during the systolic phase, a pressure wave increases the blood volume in the tissue. For this volumetric case, Equation (6) becomes

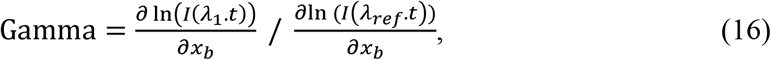

where λ_1_ = 670 or 590 nm, and λ_*ref*_ = 940 nm.

Using the transmission model given by Equation (15), the gamma value defined by Equation (16) was estimated for different sizes of aggregates ranging from 1 to 3 microns. With the RBC approximately 1.1 microns thick, a thickness of 1 micron corresponds to approximately one RBC, and a 3-micron thickness corresponds to three RBCs. To correspond to fully oxygenated arterial blood, SPO2 is taken as 1. Figure 3(a) shows that the gamma value for the wavelength pair (670 nm, 940 nm tends to decrease with the increase in the rouleau size, where the gamma value for the pair (590 nm, 940 nm) (Figure 3(b)) increases as the aggregate grows.

**Figure 3:**
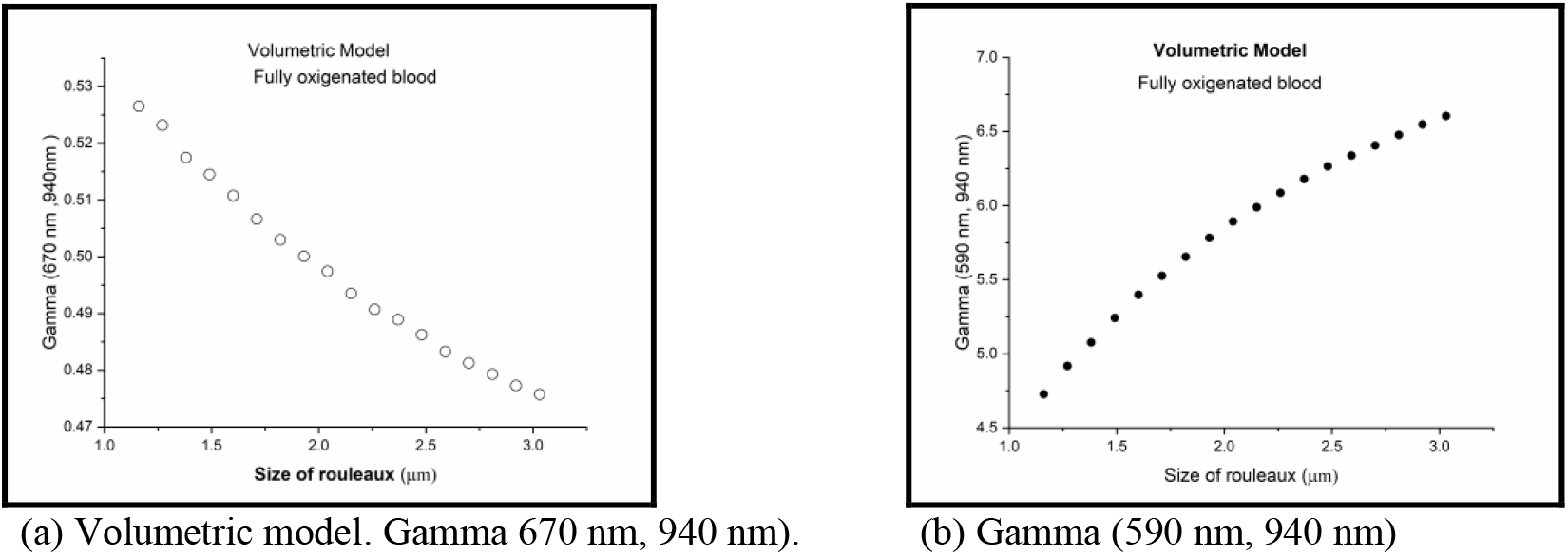

**Figure 4:**
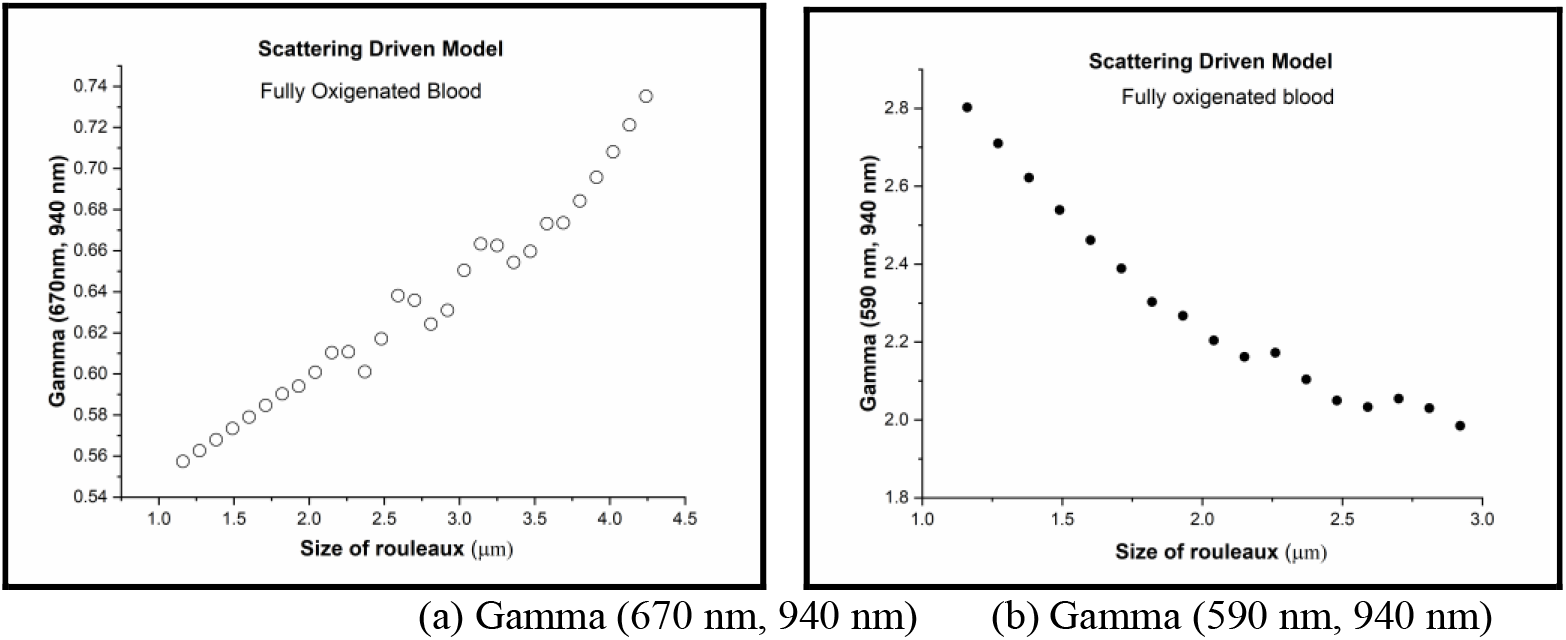
SD model.

For the SDM, the gamma values for the reflection approach (Equations (13) and (14)) and for the transmission model (Equation (15)) are calculated. The aggregation process in the SDM is the only mechanism responsible for signal changes. This process is modeled by the inflation of sphere-like scatterers, whereas the number of scatterers drops; hence, the total hematocrit remains unchanged. The particle size and concentration (c(t)) periodically fluctuate with time (t), and *μ*_*d*_(*t*) incorporates all changes that are induced by the aggregation process. From Equation (6), gamma should be written as

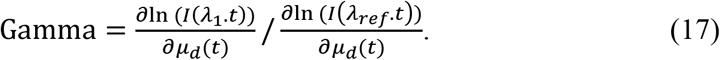

The figures below show the calculated gamma for the SDM from Equation (17) for two pairs of wavelengths, (670 nm, 940 nm) and (590 nm, 940 nm), based on the transmission model (Equation (16)).

For the reflection model (Equations (13) and (14)), the gamma values for the SDM are shown in Figure 5, (a) and (b).

**Figure 5:**
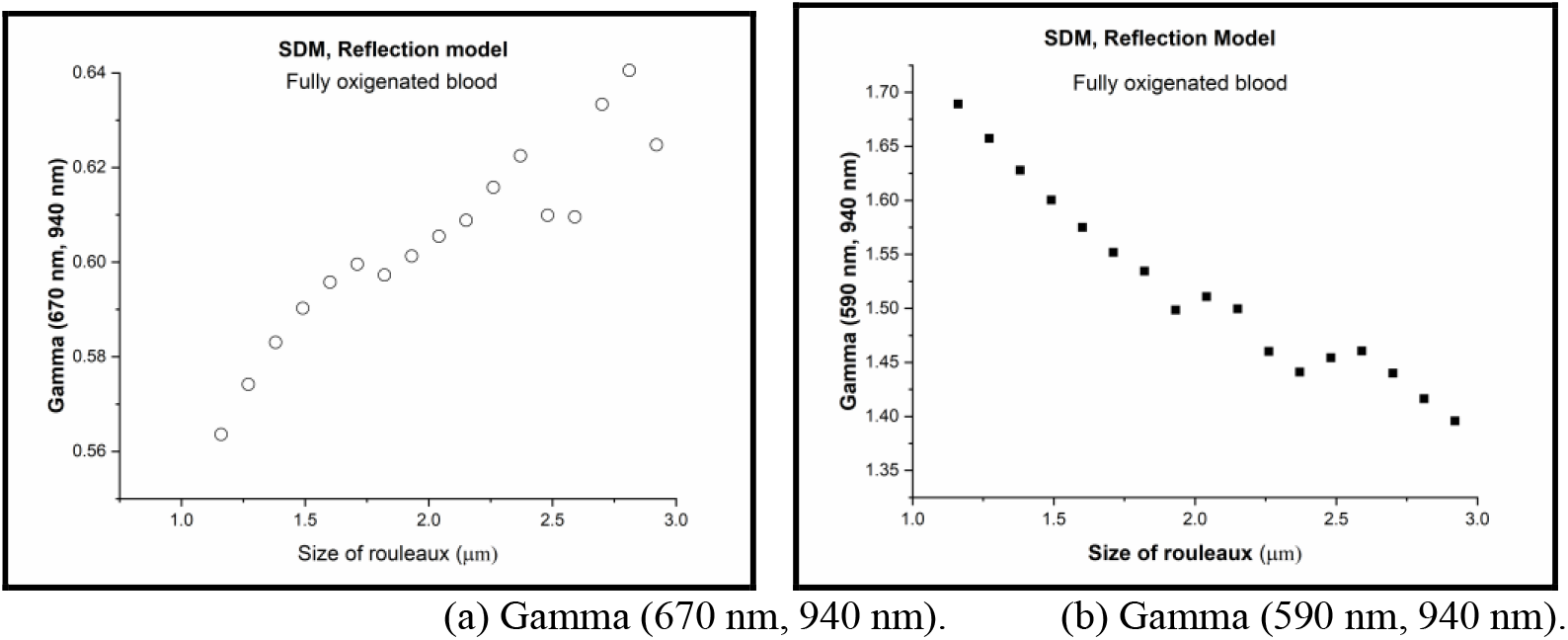
Reflection model.

With regard to the gamma dependency on the aggregate length, the reflection and transmission models for the SDM manifest similar behaviors; however, they are diametrically opposite to the volumetric model trends.

### Experiments

The measuring system (Figure 6) includes a standard reflective geometry optical scheme for measuring the PPG signal. In the experiments, two light source modules were used: matrices with LED pairs of (670 nm, 940 nm) and (590 nm, 940 nm). The distance between the light source and detector was 5 mm. To acquire PPG signals, a two-channel system based on a fully integrated chip (AFE4403, Texas Instruments) was employed; the measured signals were stored in the computer memory for further processing. An additional system module includes a pressure regulator that controls the pumping of air into a special air cuff. The cuff is cylindrical with an elastic inner membrane and a rigid outer shell. The cuff volume was minimized to increase sensitivity to volume changes. Pressure pulsations were recorded by a pressure sensor and subsequently digitized and stored in the PC memory. The experimental device had a resolution of 16 bits and a sampling rate 100 Hz.

**Figure 6:**
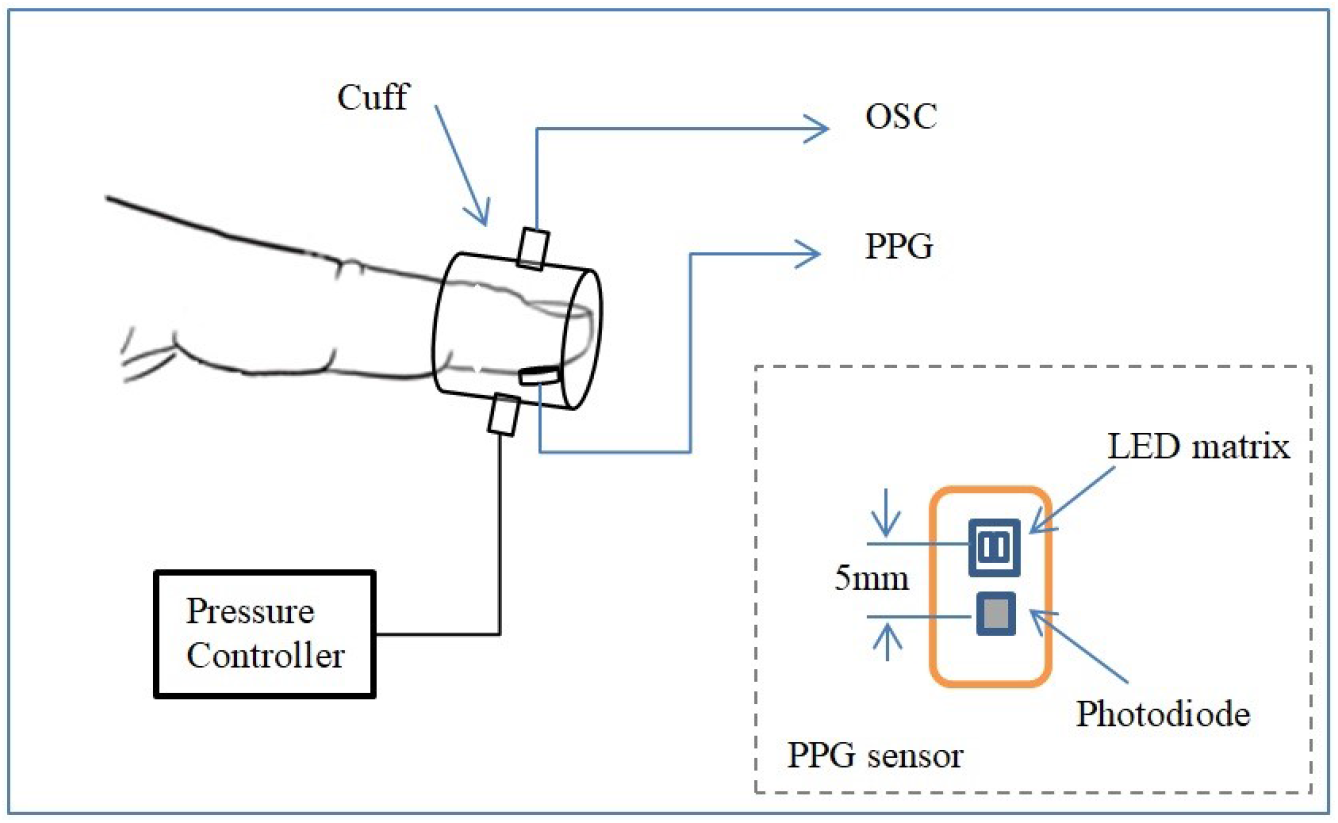
Measuring system diagram consisting of optoelectronic and pneumatic sub-systems.

Initially, a pair of LEDs (670 nm, 940 nm) was utilized, and the PPG signal from the fingertip was recorded for 1 min. At the same time, the air cuff was wrapped around the fingertip and a predetermined pressure was applied. The oscillometric signal and two PPG signals were simultaneously recorded. Then, the pressure in the air cuff was increased, and the experiment was repeated. The experiments continued until the applied pressure exceeded the systolic pressure of the subject by 40 Torrs. The entire series of experiments was repeated for another pair of LEDs (590 nm, 940 nm). Generally, the velocity of blood flowing into the fingertip is supposed to decrease with increasing pressure inside the air cuff. The velocity decrease leads to a drop in shear forces and a corresponding increase in the average length of RBC aggregates. The recorded PPG signals were processed by a separate program, and the gamma values were averaged over a 1-min period. In addition, the PPG signal waveforms at 940 nm and the oscillometric signal were recorded. Then, the amplitudes of the AC components for each type of signal and the phase shift in between were calculated.

## Results and Discussions

The comparison of the waveforms of the PPG and oscillometric signals reveals the following features. At relatively low applied pressures, the shapes of the oscillometric and PPG (940 nm) signals including the typical manifestation of the dicrotic notch are considerably similar (Figure 7(a)). Upon reaching the over-diastolic pressure at which the non-pulsatile blood flow stops, the dicrotic form disappears in the PPG signal, whereas the oscillometric signal preserves the dicrotic feature (Figure 7(b)). With further increases in pressure, a delay in the onset of the optical signal relative to the oscillometric signal is observed. This delay increases monotonically with increasing pressure (Figure 7(c)).

**Figure 7:**
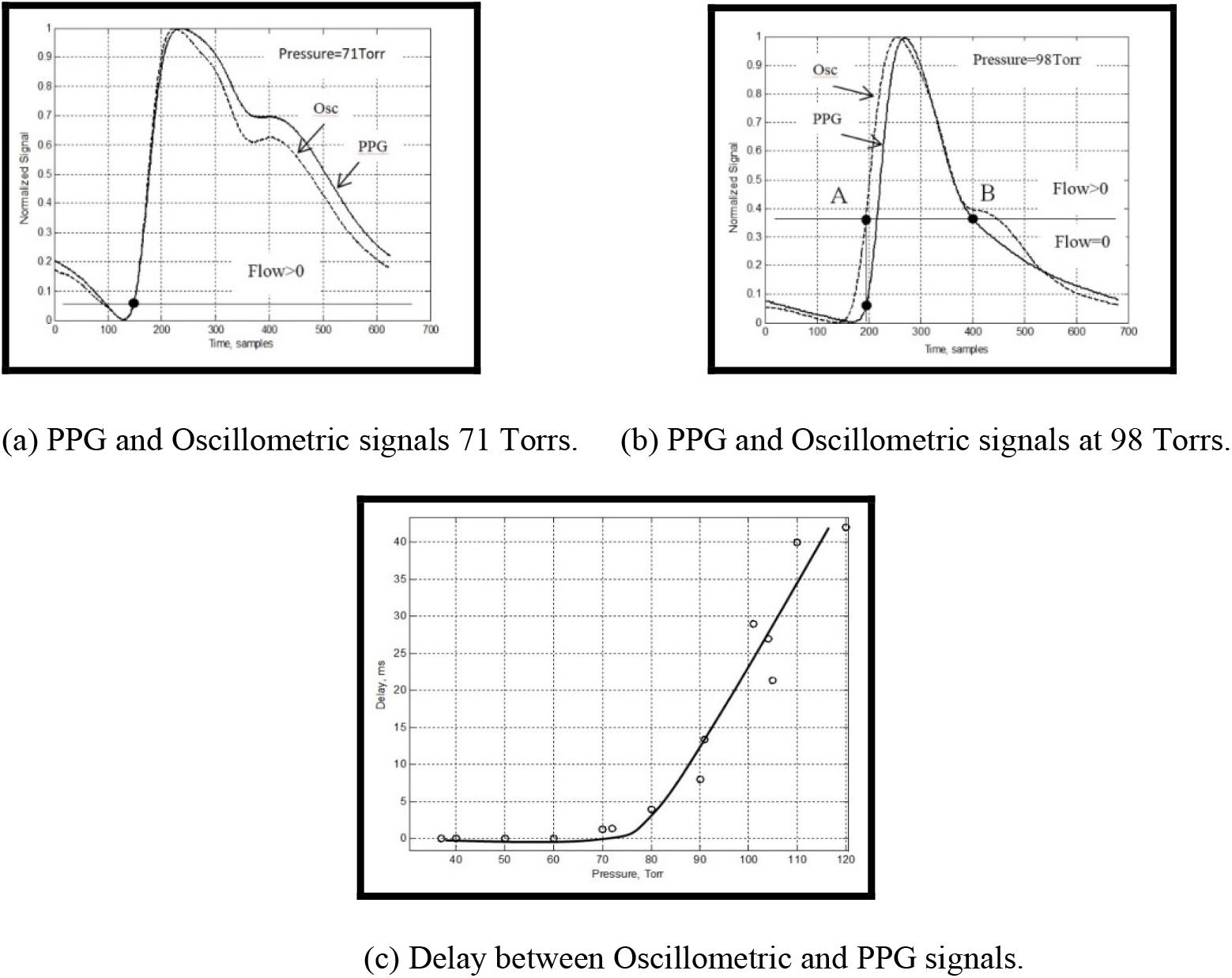

The AC value of the PPG and oscillometric signals was determined as the difference between the maximum and minimum values corresponding to the measured signals during the measurement interval. For example, at 60 Torrs, the AC value of the PPG signal was approximately13 units, and the DC value was approximately 220 units (Figure 8(a)). The amplitude of oscillometric pulsations was 0.55 Torr. Upon reaching 130 Torrs, the sufficiently significant pulsation of the oscillometric channel (0.34 Torr) and virtual disappearance of the PPG signal (Figure 8(b)) were observed. Before reaching the systolic pressure level, the dynamics of the relative decrease in the amplitude of the pulse wave oscillations for these two signals were similar (Figure 8(c)) after reaching the amplitude of the oscillometric signal.

**Figure 8:**
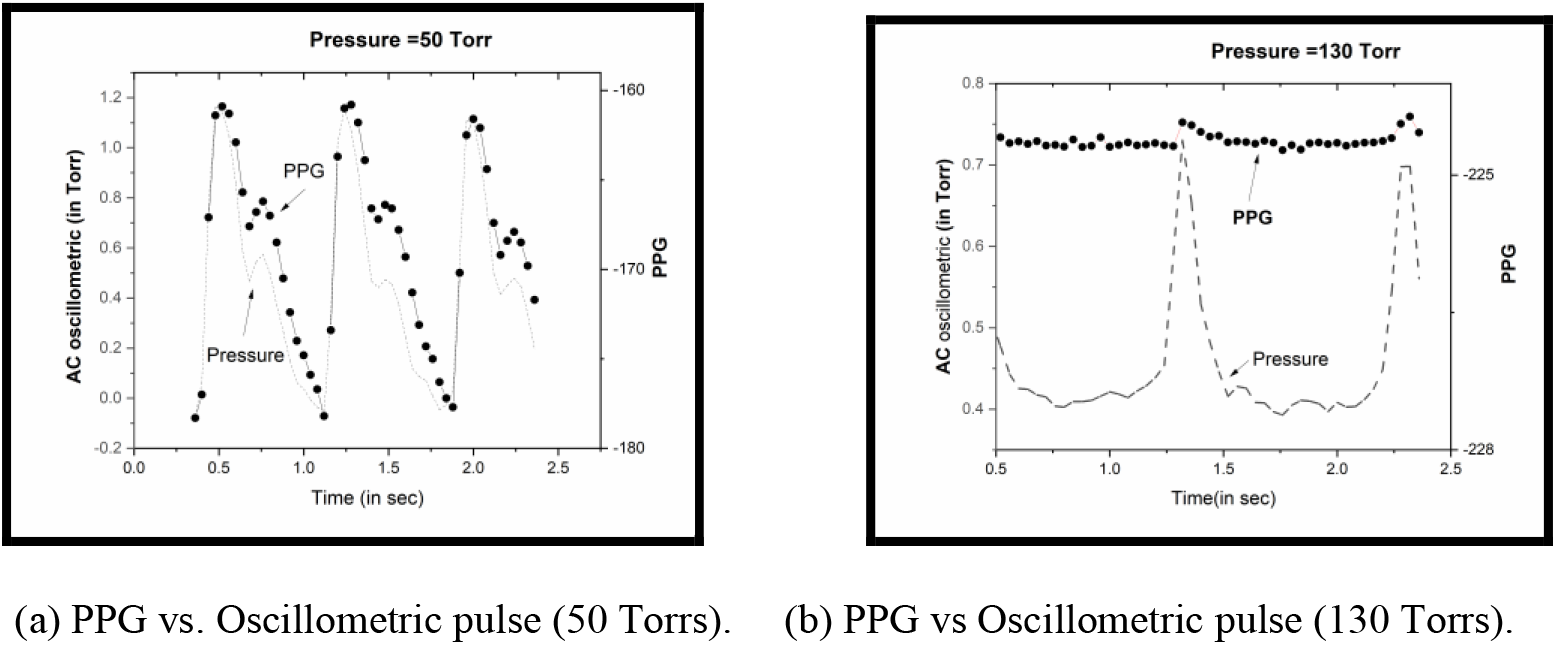
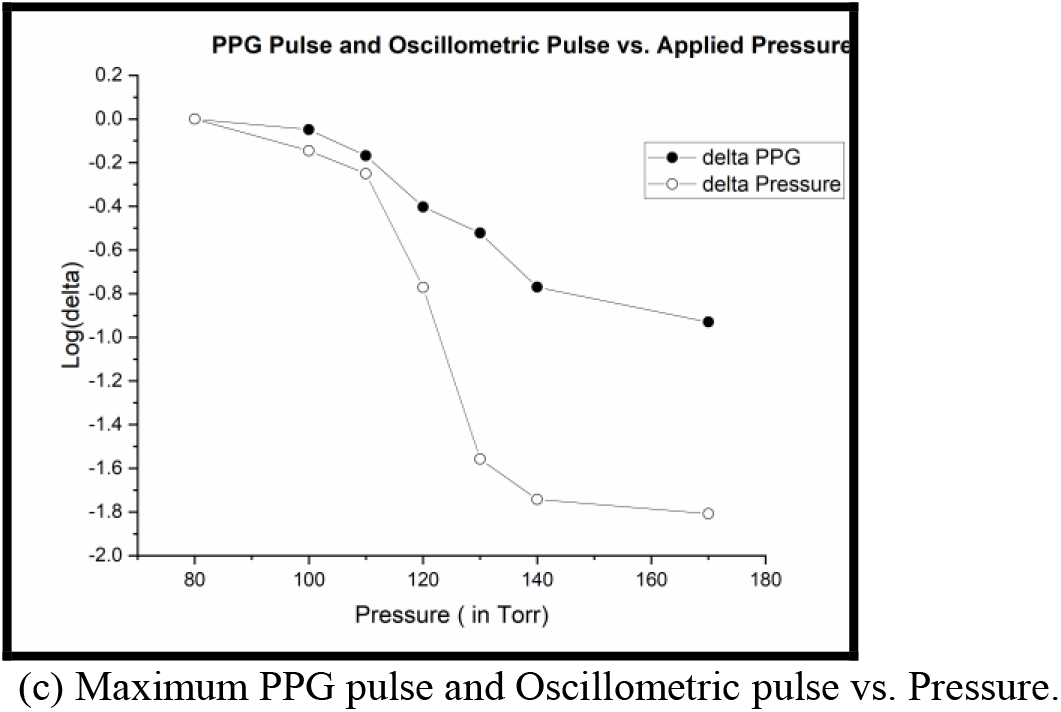

Finally, the following results show the behavior of gamma as a function of applied pressure for the two pairs of wavelengths. The experiments were repeated several times; hence, datasets from different measurements were observed. The graph shows the gamma behavior of the signal of the LED pairs, (670 nm, 940 nm) (Figure 9(a)) and (590 nm, 940 nm) (Figure 9(b)), as a function of applied pressure. The gamma value for the pair (670 nm, 940 nm) tends to increase, whereas for the pair (590 nm, 940 nm), the opposite tendency as a function of the applied pressure is observed.

**Figure 9:**
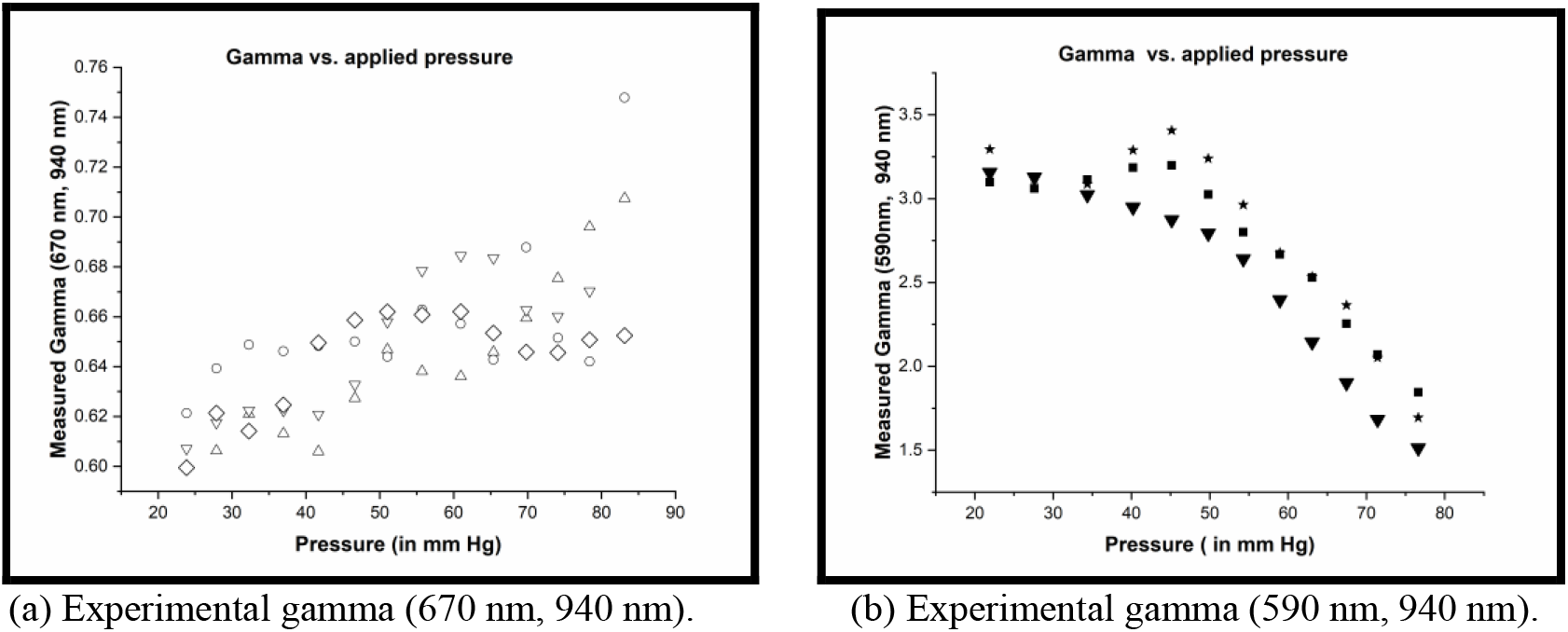

The purpose of the succeeding discussion is to compare two alternative models of the PPG signal. The volumetric model assumes that the source of the pulsation signal is the blood volume fluctuation in the vessels. According to the alternative model (i.e., SDM), the optical pulsation signal is assumed to be determined by the hemodynamic behavior of blood flow in the vessels. One of the manifestations of hemodynamics is shear force fluctuation, which leads to scattering changes in the RBCs and RBC aggregates. The first group of results is discussed to compare the simultaneously measured PPG signal (at 940 nm) and oscillometric signal associated with the volumetric behavior (Figure 7, (a)–(c)). These experiments enable the juxtaposition of the oscillometric and optical signal waveforms as a function of the applied external pressure. At low pressure levels, both waveforms exhibit similar behaviors. Upon reaching a certain threshold that approaches the diastolic pressure, the so-called dichroitic notch in the PPG signal disappears, whereas the pressure curve is preserved. According to the volumetric model, the shape of the volume change must fully correspond to the shape of the optical signal. Based on the volumetric concept, the situation in which at a certain moment the dichroitic notch disappears in the PPG but continues to be present in the oscillometric signal is difficult to explain. Another phenomenon is demonstrated in Figure 7(c). With the increase in the applied pressure, a phase lag between the pulsatile signals of the PPG and pressure wave is observed starting from a certain threshold. This lag increases with increasing pressure and is a phenomenon that is not easy to explain in terms of the volumetric model. Moreover, Figure 8(c) shows that although the PPG pulsation virtually vanishes, a considerably prominent manifestation of the oscillometric pulsation continues to be observed. This result is also inconsistent with the volumetric model. The following attempts to explain the same results based on the hemodynamic model. Within the framework of the aggregation model, the primary focus is on the behavior of the dynamics of blood flow. When the air pressure in the cuff reaches the diastolic point, the stationary blood flow stops and only the pulsating component continues. With the increase in external pressure beyond the diastolic pressure, blood flow only appears when the pulse wave exceeds the locally applied threshold pressure; this threshold is shown by the solid horizontal line in Figure 7(b). It implies that if the local arterial blood flow is measured, the delay in its appearance in relation to the pressure wave can be observed. After the expected behavior of blood flow pulsation, the observation that the PPG waveform does not correspond to the volumetric pulsation starting from a certain moment is consistent. Moreover, with the increase in the external pressure, although the amplitude of the oscillometric pulsation slightly decreases, it continues to be visible after reaching the systolic threshold, whereas the measured PPG signal disappears with the blood flow. The reason for the absence of the optical signal pulsation, whereas the volumetric pulsation continues to be observed, is difficult to explain. Thus far, the presented results indicate that the main source of the PPG signal (at least for high local pressures) is hemodynamic change and not volumetric change.

In the succeeding parts of the experiments, the correspondence between the optical model simulation of gamma and experimental results was checked. According to the proposed model, the blood flow pulsation is associated with the corresponding changes in the shear forces. These changes result in variations in the size of erythrocyte aggregates. The fluctuation in the sizes of aggregates must lead to a change in the scattering of transmitted light, as observed during the measurement. According to the proposed model, the blood flow velocity increase, which can be experimentally achieved using the applied pressure, must reduce the shear forces and increase the average size of scatterers. The model’s predictions were formulated in terms of gamma values. It has been experimentally established that for the pair (670 nm, 940 nm), the gamma value whose calculation is based on the PPG signal measured at two wavelengths does not depend on the measurement geometry and amount of blood in the tissue. This experimental fact is used in pulse oximetry. The SDM model predicts an increase in gamma for the pair (670 nm, 940 nm) and a decrease in gamma for the pair (590 nm, 940 nm) with the increase in the average aggregate length (Figures 4(a)–5(b)). Figure 9, (a) and (b), shows the experimental behavior of gamma as a function of the applied pressure measurement. For the pairs (670 nm, 940 nm) and (590 nm, 940 nm), a disagreement in the experimental result trends is observed. A strong disagreement in the gamma values for the pair (590 nm, 940 nm) is also observed.

## Conclusion

Based on the theoretical and experimental results, the following can be concluded.

- The pulsatile optical signal in the fingertip is caused by the modulation of blood flow velocity.
- The source of optical signal pulsation is associated with the modulation of the scattering of RBCs in the blood vessels.
- The change in blood scattering can be explained by the change in the average size of aggregates following the fluctuations of shear forces, which vary during the course of the pulse wave.

## Data Availability

Raw PPG data used in this study may be made available by the corresponding author upon request.

## Conflicts of Interest

The authors have no relevant financial interests in the manuscript and no potential conflicts of interest.

## Notes

### Competing Interest Statement

The authors have declared no competing interest.

